# IFNγ and antibody synergize to enhance protective immunity against *Chlamydia* dissemination and female reproductive tract reinfections

**DOI:** 10.1101/2022.08.04.502902

**Authors:** Jessica A. Gann, Priyangi A. Malaviarachchi, Wuying Du, Miguel A.B. Mercado, Lin-Xi Li

**Affiliations:** Department of Microbiology and Immunology, University of Arkansas for Medical Sciences, Little Rock, Arkansas 72205

**Keywords:** Chlamydia, IFNγ, antibody, dissemination, reinfection

## Abstract

CD4 T cell-dependent IFNγ production and antibody are the two best known effectors for protective immunity against *Chlamydia* female reproductive tract (FRT) infection. Nevertheless, mice lacking either IFNγ or B cells are capable of clearing vast majority of *Chlamydia* from the female reproductive tract (FRT), while suffering from varying degrees of disseminated infection. In this study, we investigated whether IFNγ and B cells play complimentary roles in host defense against *Chlamydia* and evaluated their relative contributions in systemic and mucosal tissues. Using mice deficient in both IFNγ and B cells (IFNγ^-/-^ x µMT), we showed that mice lacking both effectors are highly susceptible to lethal systemic *Chlamydia* dissemination. Passive transfer of immune convalescent serum, but not recombinant IFNγ, reduced bacterial burden in both systemic and mucosal tissues in IFNγ^-/-^ x µMT mice. Moreover, we observed a reduction of bacterial shedding of more than two orders of magnitude in IFNγ^-/-^ x µMT mice following both *Chlamydia muridarum* and *Chlamydia trachomatis* infections. Lastly, protective immunity against *C. muridarum* reinfection was completely abrogated in the absence of IFNγ and B cells. Our results suggest that IFNγ and B cells synergize to combat bacterial dissemination, while an IFNγ and B cell-independent mechanism exists for host resistance to *Chlamydia* in the FRT.

## Introduction

The obligate intracellular bacterium *Chlamydia* causes a variety of human and animal diseases by invading multiple mucosal tissues. Depending on the strains and serovars, *Chlamydia* may target the epithelium of the eyes, the respiratory tract, the reproductive tract, or further evade the local draining lymph nodes and cause systemic infections. *Chlamydia trachomatis* is the etiological agent for the most prevalent sexually transmitted infection with nearly 130 million cases worldwide annually (1). *C. trachomatis* infections cause major public health concerns due to the severe disease sequelae, such as pelvic inflammatory disease, ectopic pregnancy and infertility. In the past 50 years, only one Chlamydia vaccine candidate successfully proceeded to a Phase I human clinical trial (2), and there is no human Chlamydia vaccine available to date.

*Chlamydia muridarum* is a murine pathogen that shares over 98% sequence homologous with the human strain *Chlamydia trachomatis* (3). Although *Chlamydia* infections can be spontaneously resolved in immunocompetent hosts, *C. muridarum* and *C. trachomatis* can naturally ascend to the upper female reproductive tract (FRT) and cause analogous immune-mediated pathology in mouse and human. Protective immunity against *Chlamydia* is conferred primarily by CD4 cells. The importance of CD4 Th1 cells in *Chlamydia* resistance is manifested by varying degrees of defects in bacterial control in MCHII-, TCRα/β-, interleukin 12-, IFNγ-, and iNOS-deficient mice following *C. muridarum* intravaginal infection (4–7). Specifically, mice deficient in MCHII or TCRα/β exhibit long-term, high levels of *Chlamydia* shedding from the FRT, demonstrating an absolute requirement of CD4 T cells at the site of infection (4, 5). In contrast, IFNγ-deficient mice are able to clear vast majority of *C. muridarum* from the FRT, followed by a chronic phase of low-grade bacterial shedding, and a proportion of these mice succumb to disseminated infection (5, 6)ep. The distinct phenotypes in these mouse models suggest that while CD4 T cell responses are essential, host resistance to *Chlamydia* does not strictly rely on Th1-dependent IFNγ-production.

Previous studies showed that convalescent antibodies are protective against *Chlamydia* reinfections, but B cells do not appear to participate in *Chlamydia* primary clearance since the kinetics of bacterial shedding are identical in WT and B cell-deficient (µMT) mice (8, 9). More recently, we reported that both µMT and antibody-deficient mice (AID^-/-^ x µS^-/-^) experience transient systemic *Chlamydia* dissemination at the early stage of FRT infection, with concomitant increase in *Chlamydia*-specific CD4 T cell expansion in secondary lymphoid organs (10, 11). Therefore, while B cells are dispensable for bacterial clearance from the FRT, they are essential for *Chlamydia* containment in the local mucosa. Whether *Chlamydia*-specific antibodies are detrimental or beneficial in human *Chlamydia* infection remains a topic of debate. High anti-*Chlamydia* antibody titers appear to correlate with severe immunopathology in women, yet both could simply be consequences of repeated infections (12). Given the fact that causal relationships among antibody levels, immunopathology and protective immunity are difficult to establish in human infection, animal models provide valuable tools for understanding the contribution of B cells and antibodies to *Chlamydia* resistance.

Although IFNγ and antibody have long been recognized as the two major players in anti-*Chlamydia* immunity, neither seems to be absolutely required for bacterial resistance in the FRT. It is conceivable that the relatively mild phenotypes observed in IFNγ- and B cell-deficient mice were due to functional redundancy between these two effectors. To test this possibility, and also seek the potential synergy between IFNγ and B cells, we generated mice deficient in both (IFNγ^-/-^ x µMT) and examined their susceptibility to *Chlamydia* in systemic and mucosal tissues following FRT infection. The contributions of IFNγ and B cells during *Chlamydia* secondary challenge were also investigated.

## Materials and Methods

### Mice

C57BL/6 (B6), IFNγ^-/-^ (B6.129S7-*Ifng*^*tm1Ts*^/J), µMT (B6.129S2-*Ighm*^*tm1Cgn*^/J) mice were purchased from The Jackson Laboratory (Bar Harbor, ME). IFNγ^-/-^ x µMT mice (DKO mice) were generated by crossing IFNγ^-/-^ mice with µMT mice. All mice used for experiments were 6-24 weeks old, unless otherwise noted. Mice were maintained under SPF conditions and all mouse experiments were approved by University of Arkansas for Medical Sciences Institutional Animal Care and Use Committee (IACUC).

### Bacteria

*Chlamydia muridarum* strain Nigg II was purchased from ATCC (VR-123; Manassas, VA). *Chlamydia muridarum* strain Weiss and *Chlamydia trachomatis* serovar D strain UW-3/Cx were kind gifts from Dr. Richard Morrison (UAMS). *Chlamydia muridarum* strain Nigg was kind gift from Drs. Roger Rank and V. Laxmi Yeruva (UAMS). *Chlamydia* strains were propagated in McCoy cells or HeLa 229 cells, elementary bodies (EBs) purified by discontinuous density gradient centrifugation and titrated on HeLa229 cells as previously described (10).

### Chlamydia infection and enumeration

Mice were synchronized for estrus by subcutaneous injection of 2.5 mg medroxyprogesterone (Depo-provera, Greenstone, NJ) 5-7 days prior to intravaginal infection. For intravaginal infection, 1×10^5^ *C. muridarum* in SPG buffer were deposited directly into the vaginal vault using a pipet tip. For transcervical infection, 1×10^6^ *C. trachomatis* in SPG buffer were inoculated directly into the upper FRT using a NEST device as previous described (13). To enumerate bacterial shedding from the lower FRT, vaginal swabs were collected, suspended in SPG buffer and disrupted with glass beads. Inclusion forming units (IFUs) were determined by plating serial dilutions of swab samples on HeLa 229 cells, staining with anti-MOMP mAb and enumerated microscopically. To enumerate bacteria burden within tissues, intraperitoneal (IP) lavage was collected in SPG buffer, upper FRT (ovaries, oviducts, upper 1/3 of uterine horn), lower FRT (vagina, cervix and lower 1/3 of uterine horn), spleen, kidney and lung were homogenized in SPG buffer. Tissue homogenates were disrupted with glass beads, centrifuged at 500 g for 10 min, supernatants collected and serial dilutions plated on HeLa 229 cells for IFU counts.

### In vivo immune serum, recombinant cytokine and mAb treatment

Immune convalescent serum was collected from B6 mice >90 days after *C. muridarum* Nigg II intravaginal infection. Sera were pooled, sterile filtered and aliquoted before use in in vivo treatment. Each mouse received 50 µL of immune serum by IP injection once every 4 days starting one day prior to infection. For recombinant mouse IFNγ (rIFNγ) treatment, each mouse was injected IP with 1 µg rIFNγ (PeproTech) in 500 µL PBS every other day beginning one day prior to infection. Antibodies used for in vivo treatment, including anti-IFNγ (clone XMG1.2, neutralizing), anti-IL-17A (clone 17F3, neutralizing) and anti-CD4 (clone GK1.5, depleting), were obtained from BioXcell. Antibody treatment was performed by IP injection of 0.25 mg of each antibody on days -1, 1, 4, 7 after secondary challenge.

### Statistical analysis

Statistical analysis was performed with GraphPad Prism 9. Log-rank Mantel-Cox test was used for survival curves; unpaired *t* test was used for normally distributed continuous-variable comparisons; Mann-Whitney U test was used for nonparametric comparisons; Repeated measures two-way ANOVA was used to compare bacterial shedding over time.

## Results

### IFNγ and B cell double knockout mice succumb to lethal disseminated C. muridarum infection

Previous studies have demonstrated that IFNγ^-/-^ mice were highly susceptible to *Chlamydia* dissemination with mortality rates ranging from ∼30% to 100% following *C. muridarum* intravaginal infections (5, 6, 14). In contrast, B cell-deficient µMT mice experience only transient, non-lethal *Chlamydia* dissemination at the early stage of *C. muridarum* infection (10). To determine whether IFNγ and B cells elicit complementary effector functions in *Chlamydia* containment in the FRT mucosa, we generated IFNγ and B cell double deficient mice (IFNγ^-/-^ x µMT) and infected these mice intravaginally with *C. muridarum* (strain Nigg II). We observed that IFNγ^-/-^ x µMT mice succumb to infection quickly, with a mediam survival of 13 days (range 12-15 days; Fig. 1A). Although 100% of IFNγ^-/-^ mice also succumb to infection, these mice survived much longer than IFNγ^-/-^ x µMT mice, with a mediam survival of 27 days (range 15-45 days; Fig. 1A). To examine the kinetics and extent of bacterial dissemination in IFNγ^-/-^ x µMT mice, various systemic tissues from WT and IFNγ^-/-^ x µMT mice were harvested between days 4, 8 and 12-post infection and bacterial burdens measured. Although no significant difference was observed at day 4 post infection (4 dpi) (Fig. 1B), a trend of increase in bacterial burden in systemic tissues, including spleen, intraperitoneal cavity (IP lavage), lungs and kidneys, was evident in IFNγ^-/-^ x µMT mice at day 8 (Fig. 1B). By day 12, high bacterial burdens of >10^6^ *C. muridarum* were detected in both mucosal and systemic tissues of IFNγ^-/-^ x µMT mice, and these mice quickly succumb to infection after this time point (Fig. 1A and 1B). These results suggest that mice deficient in both IFNγ and B cells are significantly more susceptibility to *Chlamydia* dissemination than IFNγ or B cell single deficient mice.

**Fig. 1.**
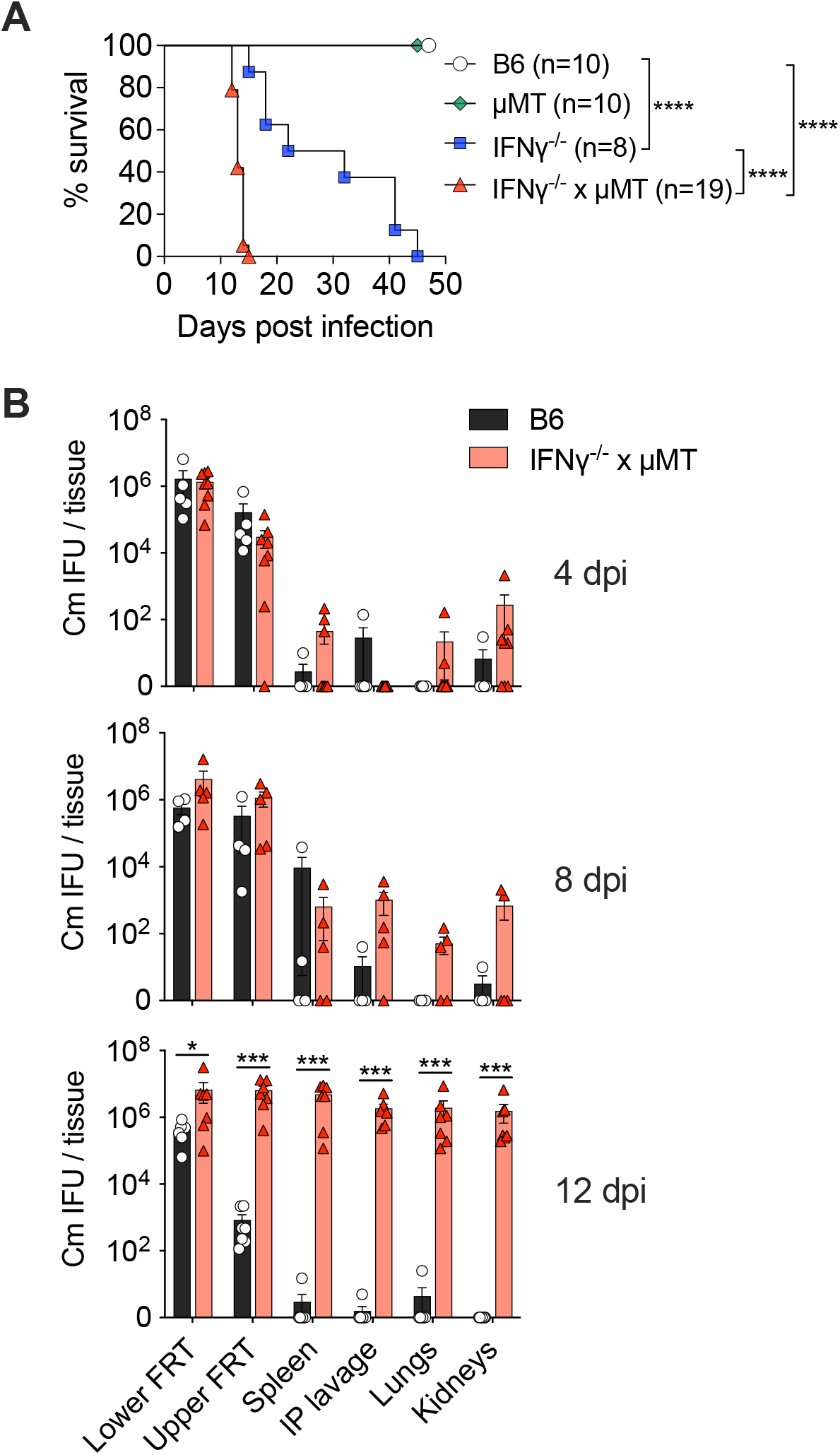
Mice deficient in both IFNγ and B cells are highly susceptible to lethal disseminated *C. muridarum* intravaginal infection. C57BL/6 (B6), IFNγ^-/-^, µMT, and IFNγ^-/-^ x µMT mice were infected intravaginally with 1×10^5^ *C. muridarum*. (A) Survival curve following infection. (B) Bacterial burdens in lower female reproductive tract (FRT), upper FRT, spleen, intraperitoneal (IP) lavage, lungs and kidneys on 4-, 8-, and 12-days post infection (dpi) as measured by IFU assay. Each data point represents an individual mouse. Data shown are pooled results of two independent experiments. Data are mean ± SEM, *p < 0.05, ***p < 0.001, ****p < 0.0001 as calculated using Log-rank test (A) and Mann-Whitney U Test (B).

### Immune serum and exogenous IFNγ partially restore C. muridarum resistance in IFNγ^-/-^ x µMT mice

During primary infection, immune serum transfer was sufficient to revert *C. muridarum* systemic dissemination in B cell deficient µMT mice (11). We therefore asked whether replenishing IFNγ^-/-^ x µMT mice with *Chlamydia*-specific antibody and/or exogenous IFNγ could restore efficient control of systemic infection in these mice. µMT, IFNγ^-/-^ and IFNγ^-/-^ x µMT mice were treated with immune convalescent serum and/or recombinant mouse IFNγ (rIFNγ) over the first 11 days of *C. muridarum* intravaginal infection, and tissues harvested at day 12 for bacterial burdens (Fig. 2A). Consistent with our previous findings, transfer of immune serum reduced bacterial burden in µMT mice in systemic tissues including spleen, IP cavity, lungs and kidneys, but not in the FRT mucosa (Fig. 2B). Likewise, replenishing circulating IFNγ, but not immune serum, to IFNγ^-/-^ mice resulted in small but significant reductions in bacterial burden in IP cavity, lungs and kidneys (Fig. 2B). Notably, in IFNγ^-/-^ x µMT mice, treatment of rIFNγ had no detectable effect on bacterial burdens in any tissue, while transfer of immune serum was able to significantly reduce bacterial counts to levels comparable to IFNγ^-/-^ mice. Lastly, treating IFNγ^-/-^ x µMT mice with rIFNγ in conjunction with immune serum had no additive effect in reducing bacteremia (Fig. 2B). Together, these data suggest that IFNγ and antibody likely elicit distinct effector functions that do not compensate for the loss of one another in host resistance to *Chlamydia* dissemination.

**Fig. 2.**
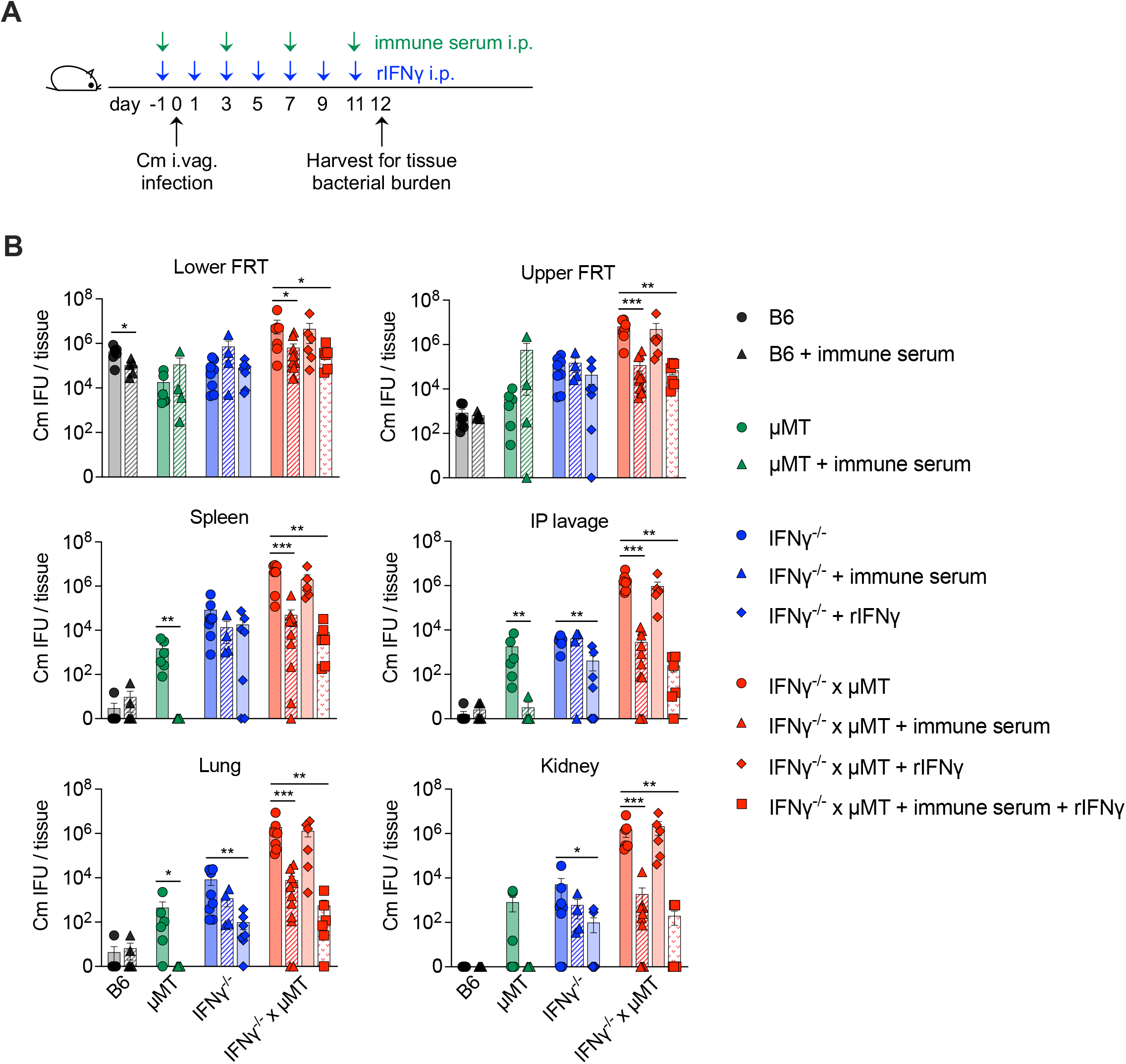
Immune serum, but not systemic IFNγ treatment, reduces tissue bacterial burden in IFNγ^-/-^ x µMT mice. B6, IFNγ^-/-^, µMT, and IFNγ^-/-^ x µMT mice were infected intravaginally with 1×10^5^ *C. muridarum*. Cohorts of mice were treated systemically with convalescent immune sera or recombinant mouse IFNγ (rIFNγ) throughout the course of infection. (A) Schematic depicting timeline for infection, immune sera and/or rIFNγ treatment, and tissue harvest. (B) Bacterial burdens of lower FRT, upper FRT, spleen, IP lavage, lungs and kidneys on day 12-post infection as measured by IFU assay. Each data point represents an individual mouse. Data shown are pooled results of three independent experiments. Data are mean ± SEM, *p < 0.05, **p < 0.01, ***p < 0.001 as calculated using Mann-Whitney U Test (B6 and µMT groups) and One-way ANOVA (IFNγ^-/-^ and IFNγ^-/-^ x µMT groups).

### Chlamydia shedding from the FRT is reduced in the absence of both IFNγ and B cells

We next determined the absolute requirement of IFNγ and B cells for reducing *Chlamydia* burdens within the FRT. It has been well documented that IFNγ^-/-^ mice are capable of clearing >99% of *C. muridarum* from the FRT while not capable of completely eradicate the bacteria (5, 6, 14). On the other hand, B cell-deficient µMT mice clear *C. muridarum* from the FRT with the same efficiency as WT controls (8, 10). Since it takes ∼5 weeks for natural resolution of *C. muridarum* from WT mice, the early death of IFNγ^-/-^ x µMT mice from disseminated infection prevented us from evaluating the full kinetics of bacterial shedding. To overcome this limitation, we infected cohorts of WT and IFNγ^-/-^ x µMT mice with three common *C. muridarum* strains (Nigg II, Weiss and Nigg) showing a spectrum of virulence (15). As suggested by survival experiments in Fig. 3A, *C. muridarum* strain Weiss exhibit slightly reduced virulence compared to *C. muridarum* Nigg II, and *C. muridarum* Nigg appeared to be the least virulent strain in this infection model. When bacterial shedding was monitored by vaginal swabs in surviving mice, significant reductions were observed in both WT and IFNγ^-/-^ x µMT mice between days 7 and 14 regardless of *C. muridarum* strain used (Fig. 3B-D). Strikingly, 5 of 12 IFNγ^-/-^ x µMT mice infected with *C. muridarum* Nigg survived at least 26 days, and bacterial burdens in these mice were more than 3 orders of magnitude lower between days 26 and 28 compared to day 7 (Fig. 3D). These observations indicated that mice are capable of reducing *C. muridarum* burdens within the FRT in the absence of both IFNγ and B cells, and alternative protective mechanisms exist for mucosal defense against *Chlamydia*.

**Fig. 3.**
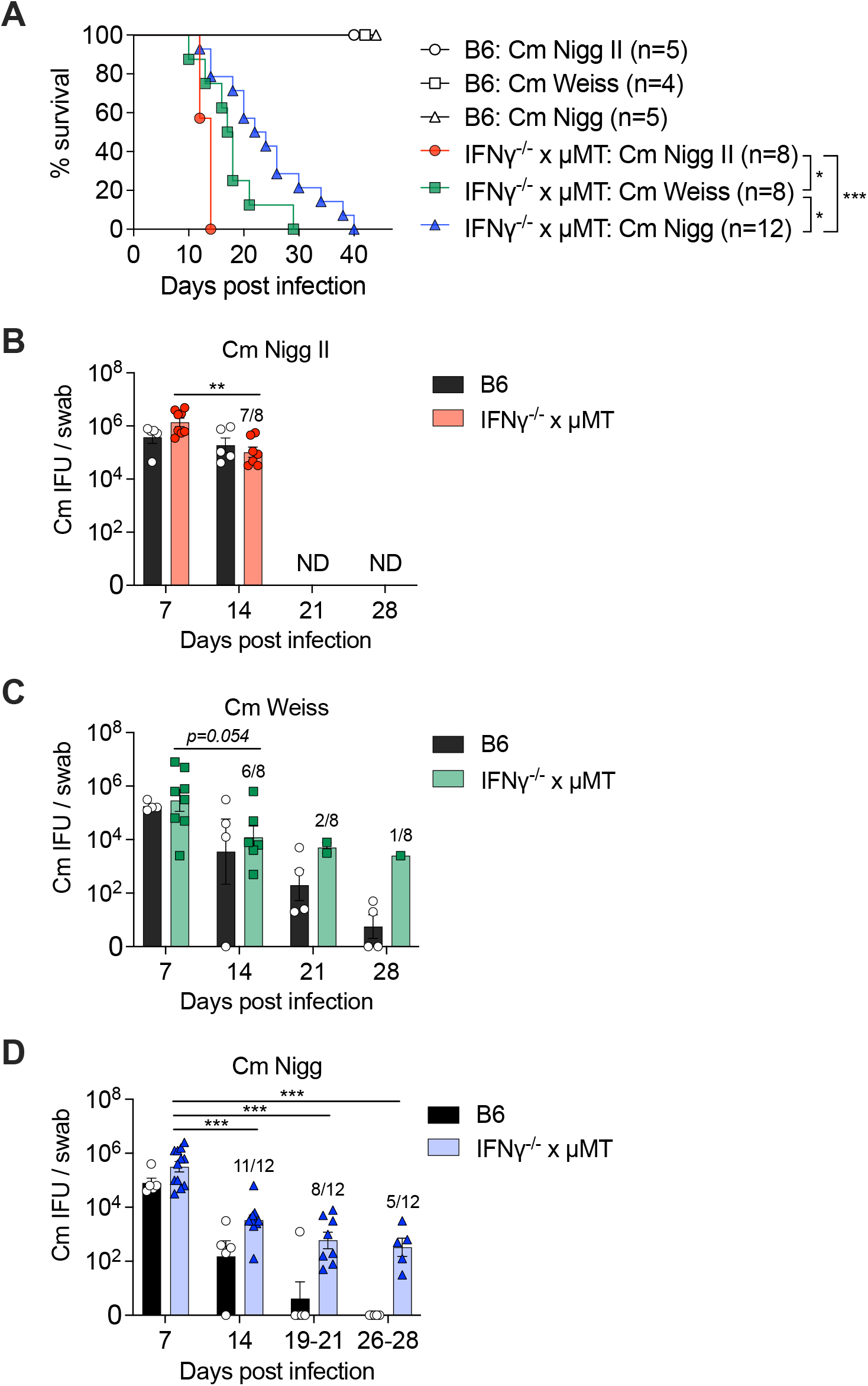
*C. muridarum* shedding from the FRT is reduced in the absence of both IFNγ and B cells. Cohorts of B6 and IFNγ^-/-^ x µMT mice were infected intravaginally with various *C. muridarum* strains (Nigg II, Weiss or Nigg) at 1×10^5^ IFU. (A) Survival curve following infection. (B-D) Bacterial shedding from the lower FRT as measured by weekly vaginal swabs and IFU assay. Numbers above the bars depicts the number of surviving mice at the time point / total number of mice in the group. Each data point represents an individual mouse. Data shown are pooled results of two independent experiments. Data are mean ± SEM, *p < 0.05, **p < 0.01, ***p < 0.001 as calculated using Log-rank test (A) and Mann-Whitney U Test (B-D). ND, not determined.

### B cells are dispensable for C. trachomatis primary clearance in mice

Intravaginally inoculation of human *C. trachomatis* in mice leads to spontaneous clearance in the absence of adaptive immunity (16). In contrast, direct deposit of *C. trachomatis* into the upper FRT requires CD4 T cells for clearance (13). Thus, intrauterine *C. trachomatis* infection provides another useful model to investigate the necessity of immune components in host resistance to *Chlamydia*. To investigate whether loss of both IFNγ and B cells affects host resistance against *C. trachomatis*, we performed transcervical infection of WT, µMT, IFNγ^-/-^ and IFNγ^-/-^ x µMT mice with 10^6^ *C. trachomatis* serovar D, and monitored bacterial shedding by vaginal swabs. WT and µMT mice quickly cleared *C. trachomatis* from FRT with very low numbers of *C. trachomatis* recovered by vaginal swabs at all time points. In contrast, IFNγ^-/-^ and IFNγ^-/-^ x µMT mice experienced prolonged bacterial shedding, and exhibited similar kinetics over the first 100 days (Fig. 4A). Although total bacterial shed from IFNγ^-/-^ x µMT mice appeared to be slightly more than IFNγ^-/-^ mice, differences between these two strains were not statistically significant (Fig. 4B). These observations supported previous findings that IFNγ is the predominant effector for *C. trachomatis* control in the mouse model, whereas B cells play a minimal role. Curiously, bacterial burdens in both IFNγ^-/-^ and IFNγ^-/-^ x µMT peaked around 14 dpi at ∼10^4 IFU and diminished to ∼10 IFU around day 80, indicating that additional effector mechanism other than IFNγ and B cells also exist for protective immunity against *C. trachomatis*.

**Fig. 4.**
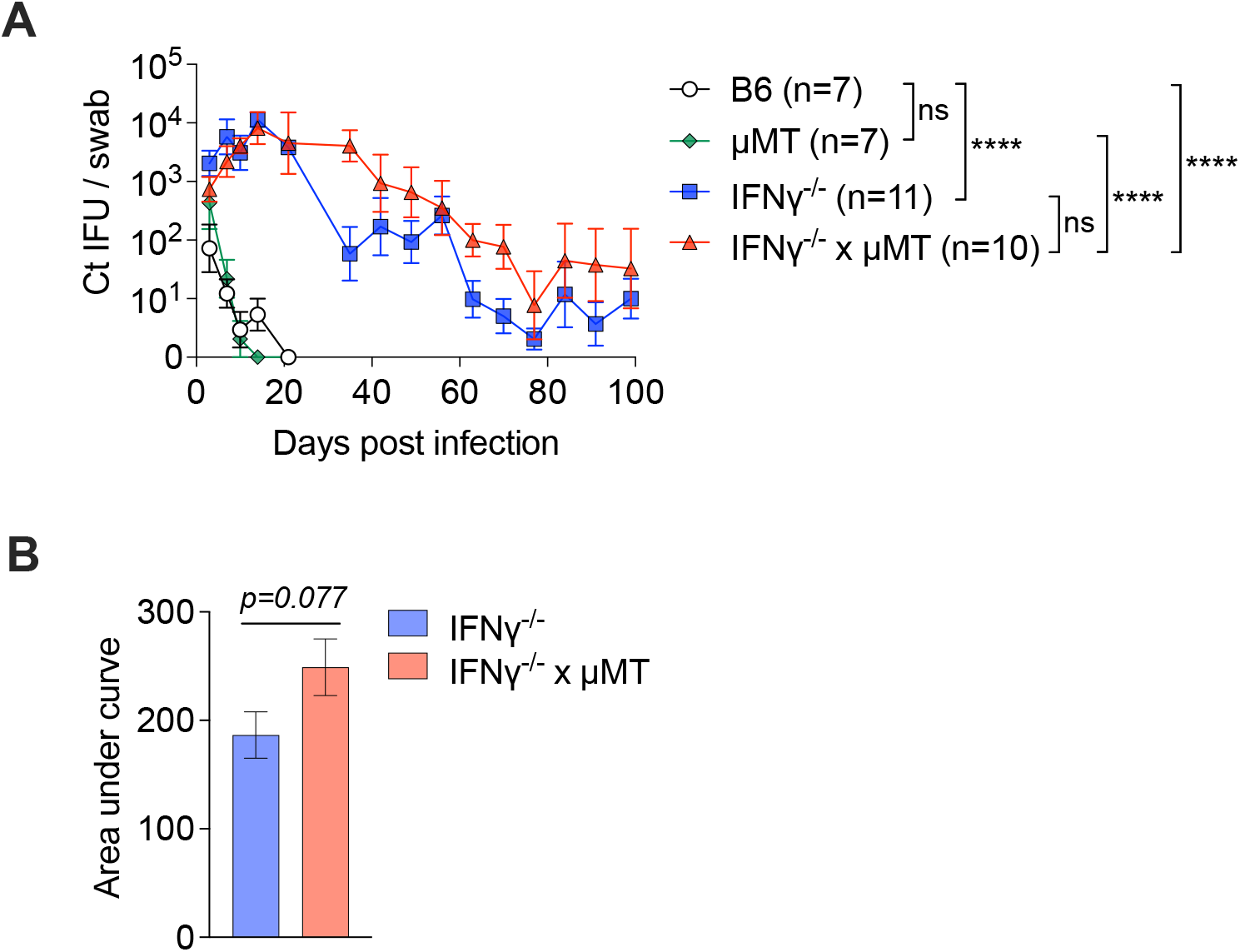
Susceptibility to *C. trachomatis* transcervical infection is not altered by B cell deficiency. B6, IFNγ^-/-^ and IFNγ^-/-^ x µMT mice were infected transcervically with 1×10^6^ *C. trachomatis* serovar D. (A) Bacterial shedding from the lower FRT were measured for the first 100 days of primary infection. (B) Area under the curve of the IFNγ^-/-^ and IFNγ^-/-^ x µMT curves in (A). Data shown are combined results of two independent experiments with 6-12 mice per group. Error bars represent the mean ± SEM, ****p < 0.0001 as calculated using Two-way ANOVA (mixed-effects analysis) (A) and un-paired t test (B).

### IFNγ and B cell double deficiency results in complete loss of protective immunity against C. muridarum secondary infection

In contrast to the absolute requirement of CD4 T cells during *Chlamydia* primary infections, either CD4 T cell or antibody can facilitate secondary clearance in mice (9). We next asked whether IFNγ and B cells play complementary roles in host resistance to *C. muridarum* reinfections, as IFNγ and B cell single deficient mice do not exhibit any major defect in secondary clearance (8, 17). To do this, we first infected µMT mice intravaginal with *C. muridarum* and allowed infections to spontaneously resolve, then rechallenged these µMT memory mice with or without antibody (Ab) treatment to neutralize IFNγ, IL-17A or deplete CD4 T cells (Fig. 5A). As shown in Fig. 5B, µMT memory mice exhibit marked protection against reinfection as bacterial burdens in isotype control treated group were >10,000 fold lower than naïve µMT mice at both day 3 and day 9. While anti-IL-17A treatment had no effect on bacterial burden in µMT memory mice, IFNγ neutralization or CD4 T cell depletion resulted in complete loss of protective immunity, and bacterial burdens in these mice were comparable to unimmunized naïve µMT controls. Additionally, neutralizing IL-17A in conjunction with IFNγ had no additive effect on bacterial burden. Taken together, we concluded that unlike in primary infection, IFNγ and B cells are the two required immune effectors for protective immunity against *C. muridarum* reinfection in mice.

**Fig. 5.**
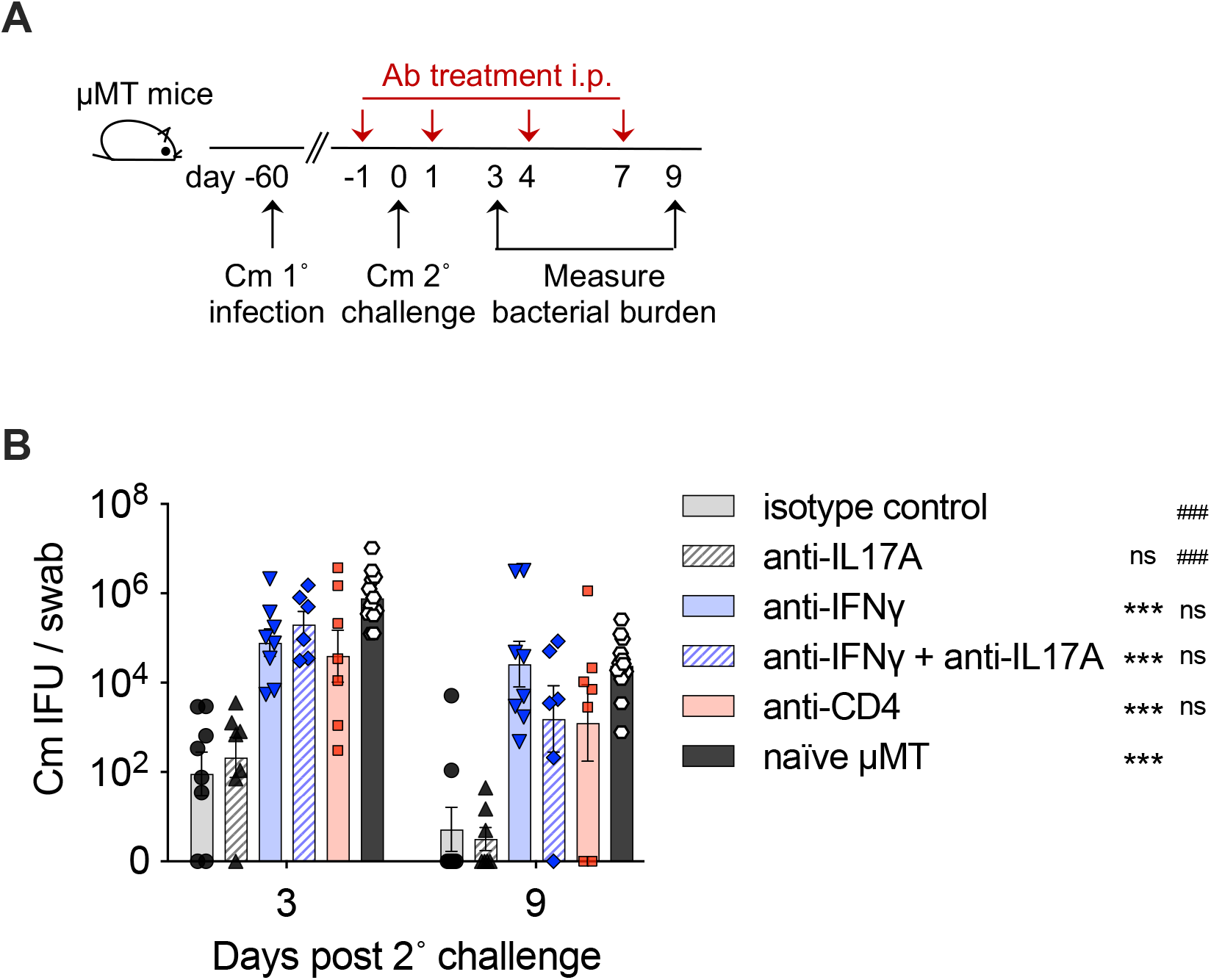
Complete loss of protective immunity against *C. muridarum* secondary infection in the absence of IFNγ and B cells. Immune µMT mice were reinfected with 1×10^5^ *C. muridarum* with or without depleting antibody treatment. (A) Schematic depicting timeline for 1° and 2° infections, antibody (Ab) treatment, and sampling. (B) Bacterial shedding from the lower FRT as measured by vaginal swabs on days 3 and 9 and IFU assay. Data shown are combined results of four independent experiments with 6-12 mice per treatment group. Each data point represents individual mouse. Error bars represent the mean ± SEM, ***p < 0.001 when compared to isotype control group, ^###^p < 0.001 when compared to naïve µMT group. Data were analyzed using Two-way ANOVA followed by multiple Mann-Whitney U Tests.

## Discussion

Understanding host defense mechanisms against *Chlamydia* is an essential step for developing a much-needed Chlamydia vaccine. To date, IFNγ and antibody are the two best known immune effectors in *Chlamydia* immunity (18). The major defects in both IFNγ- and B cell-deficient mice following *C. muridarum* intravaginal infection, however, are disseminated bacterial infections, rather than uncontrolled *Chlamydia* shedding from the FRT (5, 6, 10). It is therefore our speculation that IFNγ and antibody may compensate for the loss of one another in the FRT during *Chlamydia* infection. Using IFNγ and B cell double deficient mice (IFNγ^-/-^ x µMT), we showed definitively in this study that the major synergistic effect of IFNγ and antibody is to prevent lethal bacterial dissemination during *C. muridarum* primary infection, while additional immune effector(s) exist for *Chlamydia* control in the FRT. In contrast, deficiency in both effectors results in a complete loss of protective immunity against *C. muridarum* secondary challenge.

Following intravaginal inoculation, IFNγ^-/-^ x µMT mice exhibit a gradual increase of systemic *Chlamydia* burden over the first 12 days which leads to lethality. This phenotype closely resembles those of the *Rag2*^-/-^*γc*^-/-^ mice reported recently (14). Given that *Rag2*^-/-^*γc*^-/-^ mice are highly defective in both innate and adaptive immunity, these results suggest that IFNγ and B cells are the two immune components necessary and sufficient to control *Chlamydia* dissemination. A prior study by Poston et al. showed that *Rag1*^*-/-*^ mice succumb to *Chlamydia* dissemination following intravaginal infection with a lethal clone of *C. muridarum* Nigg (CM001). Transfer of immune serum extended the survival time of *Rag1*^*-/-*^ mice, and transfer of naïve B cells was able to completely rescued the lethal phenotype. These observations led to their conclusion that T cell-independent IFNγ (from *Rag1*^*-/-*^ mice) and B cells cooperate to prevent mortality associated with *C. muridarum* infection (19). Our study corroborates their findings and provides direct evidence that IFNγ and antibody synergize to prevent *C. muridarum* dissemination. The non-redundant roles of IFNγ and antibody is manifested by (i) partial defects (survival and bacterial burden) observed in single knock-out animals compared to IFNγ^-/-^ x µMT double deficient mice; (ii) transfer of rIFNγ, but not immune serum, reduces systemic bacteria in IFNγ^-/-^ mice; (iii) transfer of immune serum completely eradicates systemic bacteria in µMT mice, whereas the same treatment only partially reduces bacterial counts in IFNγ^-/-^ x µMT mice. It is noteworthy that replenishing rIFNγ intraperitoneally only slightly reduced *Chlamydia* burdens in IFNγ^-/-^ mice, and no reduction was detected in IFNγ^-/-^ x µMT mice (Fig. 2). Similar observation was made by Williams et al. that while IFNγ depletion in immunocompetent Nu/+ mice increase mortality and bacterial burden in the lung following MoPn (*C. muridarum*) intranasal infection, IFNγ repletion in immunocompromised Nude mice failed to confer any reproducible protection (20). The suboptimal effect of systemic IFNγ treatment suggest that the effector function of IFNγ may require a local cytokine gradient and/or cognate interaction between IFNγ-producing and -responding cells (21). Supporting this notion, IFNγ is detected in genital secretion but not in serum or spleen after intravaginal infection (ref. (22) and data not shown). In contrast to IFNγ treatment, immune serum was able to completely eradicate systemic bacteria from µMT mice. However, *C. muridarum* burdens in FRT tissues are unaffected in both WT and µMT mice. Curiously, small but statistically significant reductions in FRT bacterial burdens were observed in IFNγ^-/-^ x µMT mice after immune serum treatment, especially in the upper FRT (Fig. 2). We speculate that the hyper-inflammatory environment in IFNγ^-/-^ x µMT mice may cause tissue leakage at the FRT that facilitates diffusion of protective antibodies into the tissue without the need of transcytosis. Future experiments will be needed to test this possibility.

Examining the full time-course of infection in IFNγ^-/-^ x µMT mice was not possible due to the lethal dissemination phenotype following *C. muridarum* intravaginal infection. Using three *C. muridarum* strains (Nigg II, Weiss and Nigg) with varying levels of virulence, we showed that IFNγ^-/-^ x µMT mice were able to partially reduce bacterial burdens in the lower FRT, especially during the first 14 days of infection. It is evident that this bacterial control mechanism is IFNγ- and B cell-independent, but likely to be CD4 T-cell dependent, since T cell-deficient mice constantly shed high levels of *C. muridarum* without any reduction for at least 50 days (5, 14, 19). The time frame between days 7 to 14 also coincide with robust clonal expansion, homing and accumulation of *Chlamydia*-specific CD4 T cells in the FRT (10). In parallel to our findings, a recent study demonstrated that T-bet/Th1 cells are dispensable for primary clearance of *C. muridarum* from the FRT (23), although a compensatory effector pathway for protective immunity is yet to be discovered.

Despite the remarkable genetic similarities between *C. muridarum* and *C. trachomatis*, these two species exhibit strong host tropism during infections (24). In contrast to the results observed by Williams et al, early studies by Zhong and de la Maza showed that recombinant IFNγ was highly effective in reducing systemic bacterial burden following Ct serovar L1 intravenous infection in mice (25). The differential sensitivities of *C. muridarum* and *C. trachomatis* to murine IFNγ were later reconciled by delineating the distinct IFNγ evasion strategies used by these two strains in their respective hosts. While *C. trachomatis* is capable of salvaging indole to evade human IFNγ induced tryptophan deprivation, it is susceptible to murine IFNγ induced immunity-related GTPases, thereby quickly eliminated from IFNγ-competent mice (26–28). IFNγ^-/-^ and IFNγR^-/-^ mice manifest prolonged bacterial shedding for at least 50 days following *C. trachomatis* intravaginal inoculation (24, 29). Antibody responses in these mice were comparable to, if not higher than their WT counterparts, despite the fact that augmented Ab responses afford no protective immunity against secondary challenge (29). By comparing the kinetics of bacterial shedding in B cell-deficient mice with or without IFNγ (µMT and IFNγ^-/-^ x µMT), we showed in this current study that IFNγ is the dominant effector for *C. trachomatis* control in mice, and B cells/antibodies do not contribute to reduced *C. trachomatis* burdens at the late stage of primary infection. Therefore, much like *C. muridarum* infection in mice, a CD4 T cell-dependent, but IFNγ-independent effector must be responsible for reducing *C. trachomatis* burdens in IFNγ^-/-^ and IFNγ^-/-^ x µMT mice. Unfortunately, little information is available to date concerning alternative protective mechanisms. We argue that identification of such mechanisms is pivotal since the IFNγ-dependent host tropism of *C. muridarum* and *C. trachomatis* strongly suggests that IFNγ resistance is a natural outcome of host-pathogen coevolution, therefore an IFNγ-independent protective mechanism would be essential for the host to combat emerging IFNγ-resistant *Chlamydia* strains.

*C. muridarum* primary infections in mice generates robust natural immunity against secondary challenges, providing a valuable system for interrogating immune effector pathways essential for host protection. A major difference between protective responses to *Chlamydia* primary and secondary infections is whether antibody can confer protection in the absence of CD4 T cells (30). Recent studies by Morrison and colleagues highlighted that the effector function of antibody rely on host IFNγ production and/or presence of neutrophils (17, 31). Our data add to their findings by showing that in the absence of B cells/antibody, IFNγ is also pivotal for protection. Anti-IFNγ treatment in µMT mice results in a complete loss of protective immunity, a phenotype that mirrors the effect of anti-CD4 treatment (Fig. 5B). Thus, IFNγ likely represents the key effector cytokine produced by memory CD4 T cells to combat reinfection. An alternative, but not mutually exclusive explanation is that IFNγ produced by innate cell types is essential for inducing effective antigen presentation and/or polarization/recruitment of memory CD4 T cells. More detailed investigation is needed to dissect the precise mechanism. It is worth noting that although a protective role of Th17 cells were reported in BALB/c mouse models following *C. muridarum* intravaginal infection (32), and a shift to Th17 response was observed in mice lacking classical Th1 responses (23), we found no difference in bacterial burden following anti-IL17A treatment in the presence or absence of IFNγ, indicating that IL17A mediated signals are neither necessary nor sufficient for protective immunity against secondary infections in this setting.

The sought for immune protective mechanisms against *Chlamydia* was impeded by lack of major phenotypes in majority of gene-deficient mouse models following *Chlamydia* FRT infection (30). On the contrary, redundancy of immune effectors in protective immunity is beneficial for the hosts, which may explain why *Chlamydia* infection in human, albeit prevalent, are mostly asymptomatic and spontaneously resolved. Our study highlighted the need for further interrogation of the interplay between known immune effectors, and exploring unknown immune components beyond IFNγ and antibody. A deeper understanding of the differences between the systemic vs mucosal, and primary vs secondary immune responses to *Chlamydia* is essential for developing a protective Chlamydia vaccine in humans.

## Acknowledgements

We thank Drs. Jason Stumhofer, Tiffany Weinkopff and Lu Huang for helpful discussions. We thank Dr. Richard Morrison and Sandra Morrison for reviewing the manuscript. This study was supported by grants from the National Institutes of Health to LXL (AI139124 and GM103625).

